# “A novel female-specific circadian clock mechanism regulating metabolism”

**DOI:** 10.1101/2020.10.05.326652

**Authors:** Tsedey Mekbib, Ting-Chung Suen, Aisha Rollins-Hairston, Kiandra Smith, Ariel Armstrong, Cloe Gray, Sharon Owino, Kenkichi Baba, Julie E. Baggs, J. Christopher Ehlen, Gianluca Tosini, Jason P. DeBruyne

## Abstract

Circadian clocks enable organisms to predict and align their behaviors and physiologies to constant daily day-night environmental cycle. Because the ubiquitin ligase Siah2 has been identified as a potential regulator of circadian clock function in cultured cells, we have used Siah2-deficient mice to examine its function *in vivo*. Our experiments demonstrate a striking and unexpected sexually dimorphic effect of *Siah2* deficiency on the regulation of rhythmically expressed genes. The absence of Siah2 in females, but not in males, altered the expression of core circadian clock genes and drastically remodeled the rhythmic hepatic transcriptome. Siah2 loss, only in females, increased the expression of 100’s of genes selectively at mid-day, resulting in a >50% increase in the number of rhythmically expressed genes, and shifted the expression of 100’s of other genes from a mid-night peak, to a mid-day peak. The combined result is a near inversion of overall rhythmicity in gene expression selectively in Siah2-deficient females. This dramatic reorganization created a substantial misalignment between rhythmic liver functions and feeding/behavioral rhythms, and consequently disrupted daily patterns of lipid/lipoprotein metabolism and metabolic responses to high-fat diet. Collectively, our results suggest that Siah2 is part of a female-specific circadian mechanism important for maintaining metabolic homeostasis and may play a key role in establishing sexual dimorphisms in metabolism, and broadly reveal that circadian clocks may drive rhythms using novel sex-specific transcriptional pathways.

## Introduction

Circadian rhythms in physiology and behavior are driven by transcriptional feed-back loop timing mechanism that drives ~24hr rhythms in expression of 1,000’s of target genes (1–3) thoughout the body. How circadian clocks drives these rhythms is thought to be due to largely similar transcriptional pathways and mechanisms in males and females, although some rhythms are modulated by sex and growth hormones. Disruption of these rhythms, or more commonly, misalignent of these rhythms with the environmental day-night cycle or within the organism causes a wide-range of health consequences, including metabolic dysfunction and obesity (4–6).

Siah2 is a Ring-type E3 ligase whose role in regulating the hypoxia pathway (7) and tumorogenesis is well known (8, 9). We have recently reported evidence suggesting that Siah2 is also a regulator of circadian clock function (10). We identified Siah2 in a screen for ubiquitin ligases that mediate degradation of RevErbα/β, heme-sensitive transcriptional repressors that regulate circadian rhythms and lipid metabolism (11–17). Suppressing *Siah2* expression in a cellular clock model U2OS cells both altered RevErbα stability and lengthened periodicity, thus directly implicating Siah2 in the regulation of circadian rhythms (10). To explore this possibility further, we have now examined the effect of Siah2 deletion on clock function in a mouse model (18). Here we present data that reveal Siah2 is a component of unexpectedly female-specific transcriptional mechanisms that are essential for the proper rhythmic control of gene expression in the liver. We also show that disrupting this mechanism substantially impairs the circadian regulation of lipid and cholesterol metabolism selectively in females and weakens their resistance to diet-induced obesity, suggesting sex-specific circadian mechanisms may contribute broadly to differences in male and female physiology.

## Results

REVERBα protein levels reach peak abundance levels during the daytime in most tissues, and in the liver, they reach peak levels around Zeitgeber time (ZT) 9-10, which is ~2-3 hours before lights go out (lights off = ZT12; Preitner et al., 2002). Since REVERBα functions a transcriptional repressor, we first asked if there were detectable changes in gene expression in livers harvested from Siah2 KO mice just after REVERBα peak levels using a QuantSeq 3’mRNA counting approach. Intriguingly, we did not detect any significant changes in expression unless we analyzed the effects of SIAH2 loss within each sex. These analyses revealed that Siah2 loss altered the levels of ~6% of detected transcripts in livers of female mice, but less than 0.1% of transcripts detected in males (Fig 1A-B; see also Supplemental Dataset 1). Of note, the transcripts altered by Siah2 loss in females appear to be enriched by >2-fold for genes that are rhythmically expressed in livers (Fig 1C), suggesting a hypothesis that Siah2 loss may impact rhythms in females. In line with this, gene ontongeny predicted that Siah2 function may regulate a number of genes that regulate circadian rhythmicity in females, including *Per3* (Fig 1D, these genes are highlighted in Supplemental Dataset 1) in addition to genes involved in processes controlled by the clock (i.e. redox, lipid metabolism). Overall these data raised the possibility that Siah2 may have a female-specific role in regulating circadian function in the liver.

**Fig. 1.**
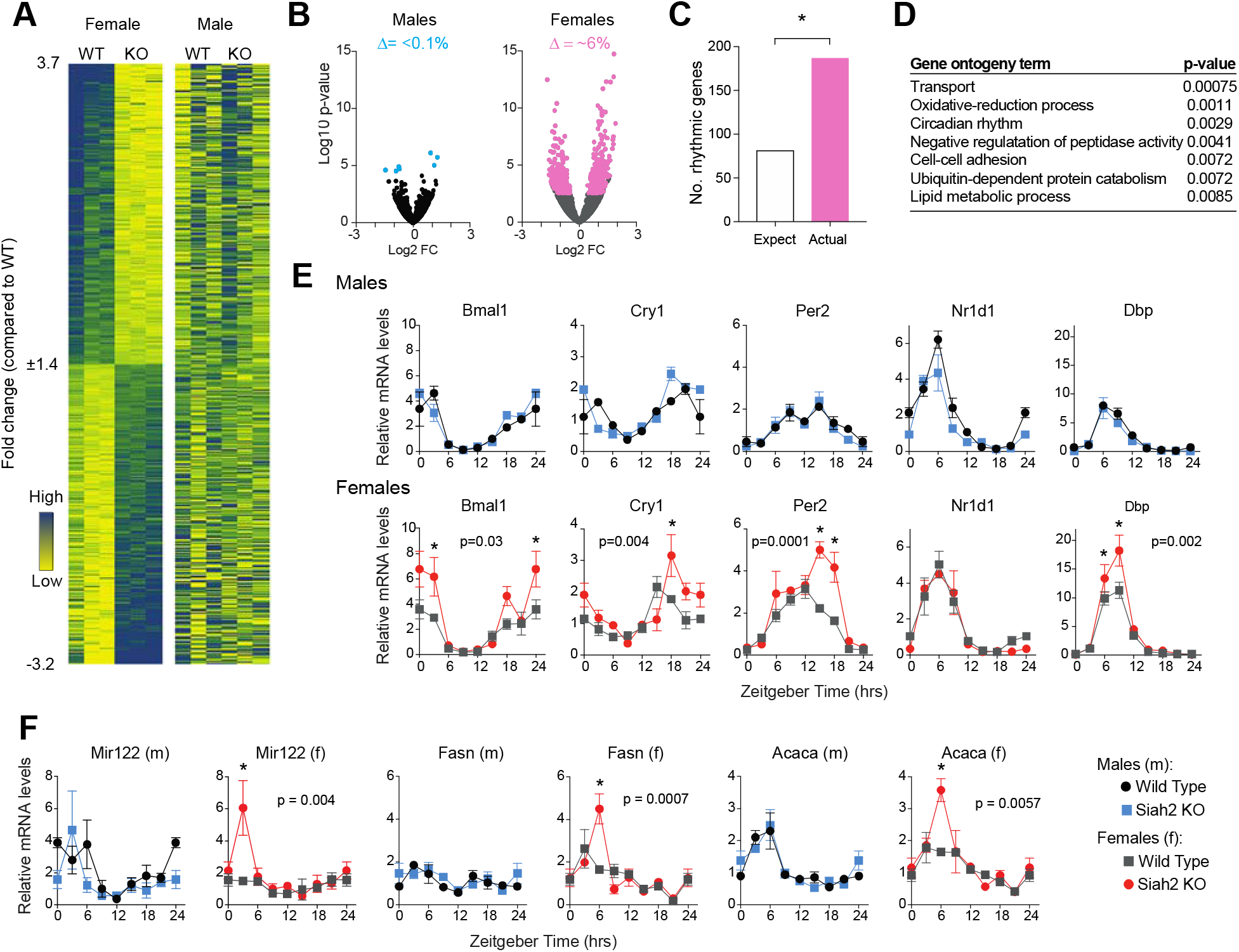
Siah2 loss preferentially alteres daily gene expression in females. **A**. Heatmaps depicting the expression of the genes affected by Siah2 loss in female livers harvested at ZT10 in all four groups. FDR corrected p-value <0.05 was used as a cutoff. Each gene is aligned across genotype (WT = wild type, KO = Siah2 KO) and sex. **B**. Volcano plots comparing hepatic gene expression changes within sex (n=3/group). Colored transcripts are signifcantly changed by Siah2 loss in male (blue) or females (pink). **C**. Enrichment of rhythmcially expressed genes among those altered by Siah2 loss in females, using 16% as the expectation for rhythmic genes (http://circadb.hogeneschlab.org/mouse; Zhang et al., 2014), * = p<0.0001 Fisher’s exact test. **D**. DAVID gene ontogeny of the rhythmic genes altered by Siah2 loss in females. Also see Supplemental Dataset 1. **E-F** Quantitative RT-PCR profiles of clock and REVERBα target gene mRNAs in liver (mean +/− sem, n=3 livers per genotype, sex and time.). The Zeitgeber time (ZT) 0 point is double plotted at ZT24 for clarity. ZT0-12 = lights on, ZT 12-24 = lights off. P-values shown reflect significant time x genotype interactions (two-way ANOVA). * = p<0.05 between genotypes at individual timepoints (Sidak’s multiple comparison test).

To explore this possibility, we examined the expression of five core rhythmic genes (*Bmal1, Cry1, Per2, Nr1d1/RevErbα* and *Dbp*) (1–3) and several REV-ERBα transcriptional targets (11, 12, 17, 19) in the livers of both male and female Siah2-deficient and wild type mice. Expression was examined at 3-hour intervals throughout a 24-hour period. We found a striking difference in the effect of Siah2 gene deletion between female and male mice (Fig1E). In male, loss of Siah2 has no detectable effect on the expression of any of these genes. In female mice, however, Siah2 loss increased the peak expression of all but one (*RevErbα*) of the core clock genes (Fig 1E) and delayed the expression of the repressors *Per2* and *Cry1* (Fig 1E). Peak expression of several REV-ERBα targets was also increased by Siah2 loss in females (Fig 1F). The effects were not limited to a specific time of day– Siah2 loss increased and/or delayed peak expression of these genes regardless of the time of day each is normally maximally expressed, suggesting that the circadian rhythm amplitude may be broadly enhanced in female livers without SIAH2. These results are somewhat different to our previous results in U2OS cells (which are female; Pontén and Saksela, 1967) where *Siah2* knockdown blunted expression of REVERBα targets coincident with altered REVERBα protein turnover (10). However, in the liver, we were surprised to find that SIAH2 loss had little effect on rhythms of REVERBα protein abundance rhythms of either sex (Suppl. Fig S1), possibly owing to tissue-specific compensation by other ubiquitin ligases (21–23). Thus, it is not clear if these effects of SIAH2 loss are through alterations in REVERBα function, though female-specific regulation of REVERBα is not expected given its role in circadian regulation. Nonetheless, the effect on gene expression in female livers suggests that Siah2 broadly regulates the amplitude and timing dynamics of circadian gene expression *in vivo*, but only in females.

We next examined the effect of Siah2 loss on the entire hepatic transcriptome via RNAseq on livers harvested around the clock. For this experiment, we chose to focus on a design that sought to strike a balance between number of animals, time resolution and comparing effects in male and females. We decided to pool RNAs extracted from livers of each three mice of sex and genotype collected at 3 hour intervals across a single day. While pooling makes it difficult to detect differences at any single timepoint like those discussed above, we expected this design would allow us to identify changes in the patterns of daily expression, reflected by substantial differences across multiple timepoints among the four groups, while simultaneously buffering against variability between individuals.

Remarkably, results from this experiment revealed that Siah2 loss in females drastically reorganized the timing of a rhythmic gene expression in livers. In both wild type males and females, most rhythmically expressed genes peaked during the night, with a population mean vector of ZT ~20-21 (Fig 2B). In Siah2 KO males, the global rhythmicity pattern appears mildly shifted (~2 hours) towards dawn. In females however, Siah2 loss shifted the the global expression profile by ~9 hours (Fig 2B), with the vast majority of rhythmic genes peaking during the daytime instead of the night. This near inversion of rhythms in female Siah2 KO livers strongly suggests Siah2 plays a central role in organizing circadian rhythms in gene expression, predominantly in females.

**Fig. 2.**
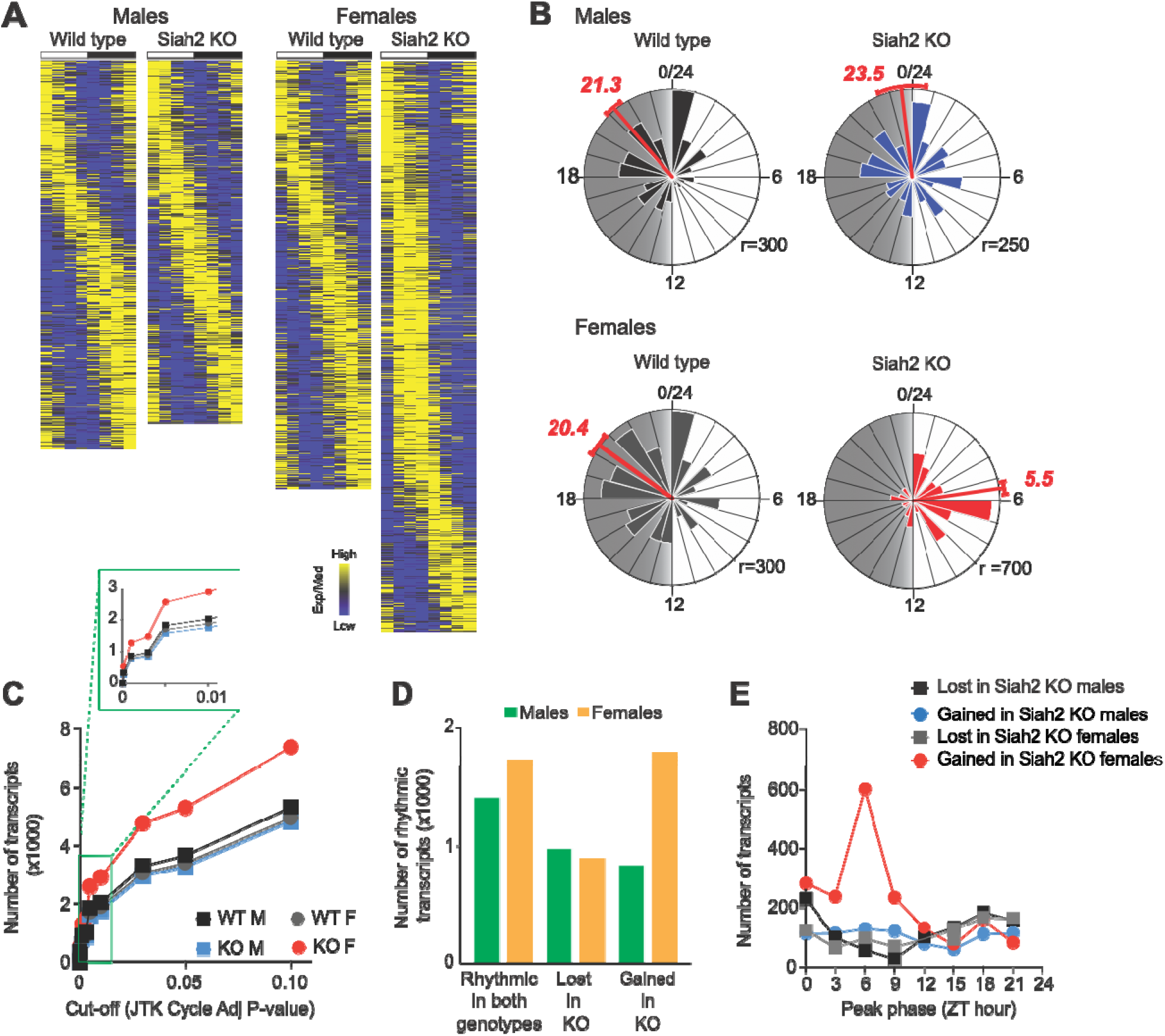
Siah2 loss selectively remodels the female circadian transcriptome towards the daytime. (**A**) Heatmaps of expression profiles of the rhythmically expressed genes (defined in Supplemental Materials and Methods), sorted by peak expression timing/phase for each group independently. White/black bars indicate the light-dark cycle. (**B**) Raleigh plots (circular histograms) of expression peak timing for all rhythmically expressed genes in each group. The numbers in red depict the average peak-time (in ZT hours) indicated by vector analysis (red line, +/− 95% CI). Gray shading = nighttime, r = radius in number of genes. **C.** Numbers of ‘rhythmically’ expressed genes in each group at various statistical cut-offs. (**D**) Comparisons of Siah2-induced changes to rhythmicity of genes in each sex. (**E**) Frequency distribution of expression peak timing, across the day, of genes that gained or lost rhythmicity in Siah2-deficient (*Siah2* KO) livers in both sexes.

This female-specific reorganization of the rhythmic transcriptome is the consequence of two principal changes. First, Siah2 loss in females resulted in a dramatic net increase (ca. 50-65%) in the number of rhythmically expressed transcripts depending on how we defined ‘rhythmicity’ (Fig 2C; see also Suppl. Fig S2 and Supplemental Methods). This increase was due to a much larger population of genes that gained rhythmicity in Siah2-deficient females over genes that lost rhythmicity (Fig 2D). In contrast, Siah2-deficiency in males caused a small net decrease (ca. 7%) in the overall number of rhythmic genes. Strikingly, most of the genes that gained rhythmicity in Siah2 KO females did so with peak expression strongly clustered around mid-day (ZT6, Fig 2E), accounting for part of the overall shift in overall timing from night time to daytime. In contrast, any changes in expression in males lacked phase clustering (Fig 2E). The large bias in the timing of expression of genes in females suggests that Siah2 regulates a *female-specific* circadian transcriptional mechanism that may prevent the expression of a large number of genes specifically around mid-day.

Second, Siah2 loss caused large shifts in when rhythmic genes peak, predominantly altering expression of nocturnally expressed genes. Examination of the patterns of genes that were rhythmically expressed in both genotypes revealed that Siah2 loss changed the time of peak expression levels of individual genes in both sexes (Fig 3A). In males, Siah2-deficiency shifted expression of genes that are expressed at all times of day, and did not alter the liver’s overall gene expression profile (Fig 3A-B). In surprising contrast, Siah2 loss in females predominantly shifted genes expressed during the night in wild type livers to peak during the day in Siah2 KO livers (Fig 3A-B). The result is a dramatic increase in the number of genes expressed during the day and reduction in the number of genes expressed during the night. Since these genes are rhythmically expressed in both genotypes, this change is the result in a phase shift in the timing of expression, in many cases by 12 hours. This is exemplified in genes that were shifted by 6-hours or more by Siah2 loss (Fig 3C) that demonstrate that genes normally peaking around midnight (ZT18) in wild type mice, shifted their expression to the complete opposite time of day. Surveying a number of individual gene profiles suggests that Siah2 loss in female livers both increased expression of genes during the day, perhaps sharing a common origin as the genes that become rhythmic in Siah2 KO females described above, and suppressed their expression during the night (Fig 3D), implicating Siah2 in different mechanisms at opposite times of day. While this shift in expression impacted 100’s of transcription factors (Fig 3D, see also Supplemental Dataset 2), expression of core circadian clock genes was not drastically shifted in female Siah2 KO liver (Fig 1E, Fig 3E). Also, behavioral rhythms locomotor acitivty and feeding behavior driven by the central circadian clock in the SCN rhythms were not affected by Siah2 loss in either sex (Fig. 3F-G). Notably, the circadian transcriptome reorganization appears to be specific to the ‘rhythmicity’ of gene expression, as we found only a few genes whose expression was altered by Siah2 loss, independent of time-of-day (Suppl. Fig S3A-B). Combined, these result suggest Siah2 may regulate two distinct mechanisms in the liver that seem to be essential in determining which and when many output genes are expressed in a circadian pattern, some of which may feedback into the clock to regulate its amplitude. These data also indicate that these unknown Siah2-dependent mechanisms are only present in females, representing a major departure from circadian regulation in males.

**Fig. 3.**
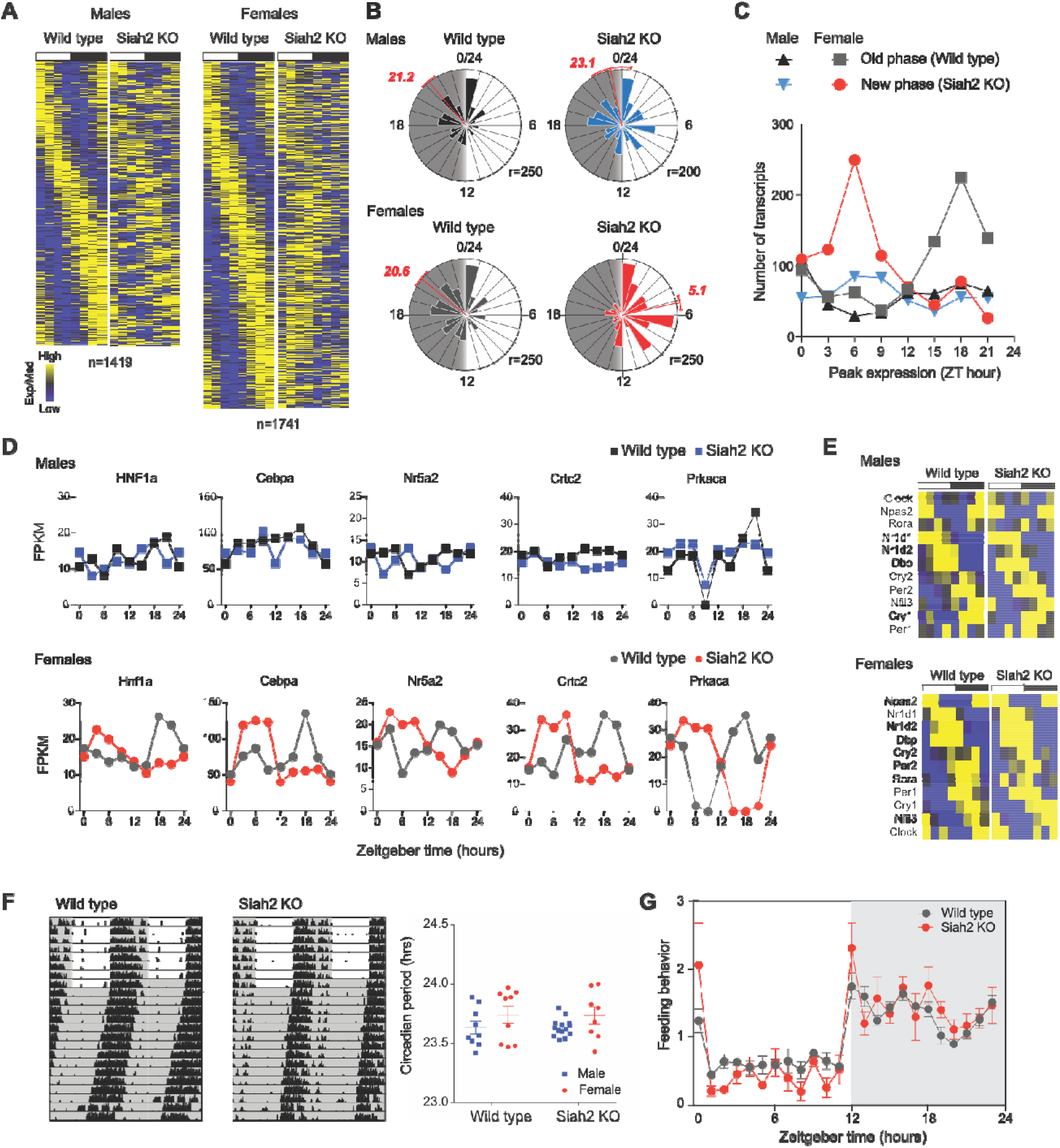
Siah2 loss in females shifts rhythmic gene expression profiles from night to day. (**A**) Heatmaps of genes that were rhythmically expressed in both genotypes; genes are aligned across genotypes (but not across sex). (**B**) Raleigh plots of the peak expression phase for the genes in A. Plotted as decribed for Fig 2B. (**C**) Frequency distributions of peak phase across time of day for genes rhythmically expressed in both genotypes but shifted by more than 6 hours between genotypes. See also **Fig. S1** and **Supplemental Datasets 2-3**. (**D**) Individual examples gene expression profiles of genes across all four groups. (**E**) Heatmaps depicting similar rhythmic expression of core circadian clock genes across all four groups. (**F**) Representative double-plotted actograms of wheel running behavior in wild type or *Siah2* KO mice. The first 8 days of the recordings (indicated by the alternating white/gray shading) were done on a 12:12LD cycle, followed by constant darkness (solid gray shading). Individual and mean +/− SEM data obtained from mice of both genotypes and sexes are shown on the right, combined from animals of both sexes and genotypes run in two independent experiments (n = 9 wild type males, 9 wild type females, 14 Siah2 KO males, and 8 Siah2 KO females). Siah2 loss did not significantly alter the behavioral circadian periods within either sex (p>0.9, two-tailed t-test, effect sizes of either genotype was than 0.006 hours). **(G)** Diurnal feeding behavior of wild type (n=5) and *Siah2* KO (n=4) female mice on normal chow. Each point is the mean of the average 8-day profile produced by each animal, and the error bars (SEM) indicate the inter-animal variability at each time point. Two-way ANOVA did not detect a significant interaction between genotype and time that would indicate a change in the daily feeding pattern (p = 0.4751 for interaction, F (23, 161) = 0.9947).

We next started exploring if other known sexually dimorphic mechanisms could be provide clues to a possible mechanism. The *Siah2* gene is not X-linked and confirmed that its expression is not different between male and female livers (Suppl. Fig S3C). We also found very little overlap in the expression of genes impacted by Siah2 loss between males and females (Fig 4A). Siah2 KO mice breed normally as homozygotes, and have normal levels of sex-hormones (Suppl. Fig S3D-E) suggesting they have at least functionally effective female reproductive endocrinology. Similarly, there was little overlap in genes altered by Siah2 loss with genes that are regulated by estrogen (Suppl. Fig S3F) or dimorphic growth hormone signaling (Suppl. Fig S3G-H; see also Supplemental Datasets 2-5). Overall, the circadian transcriptome changes in female Siah2 KO mice do not appear to be associated with alterations in the predominant hormonal mechanisms that underlie many sexual dimorphisms, raising the possibility that Siah2 may act on novel female-specific pathways to regulate rhythmic gene expression.

**Fig. 4.**
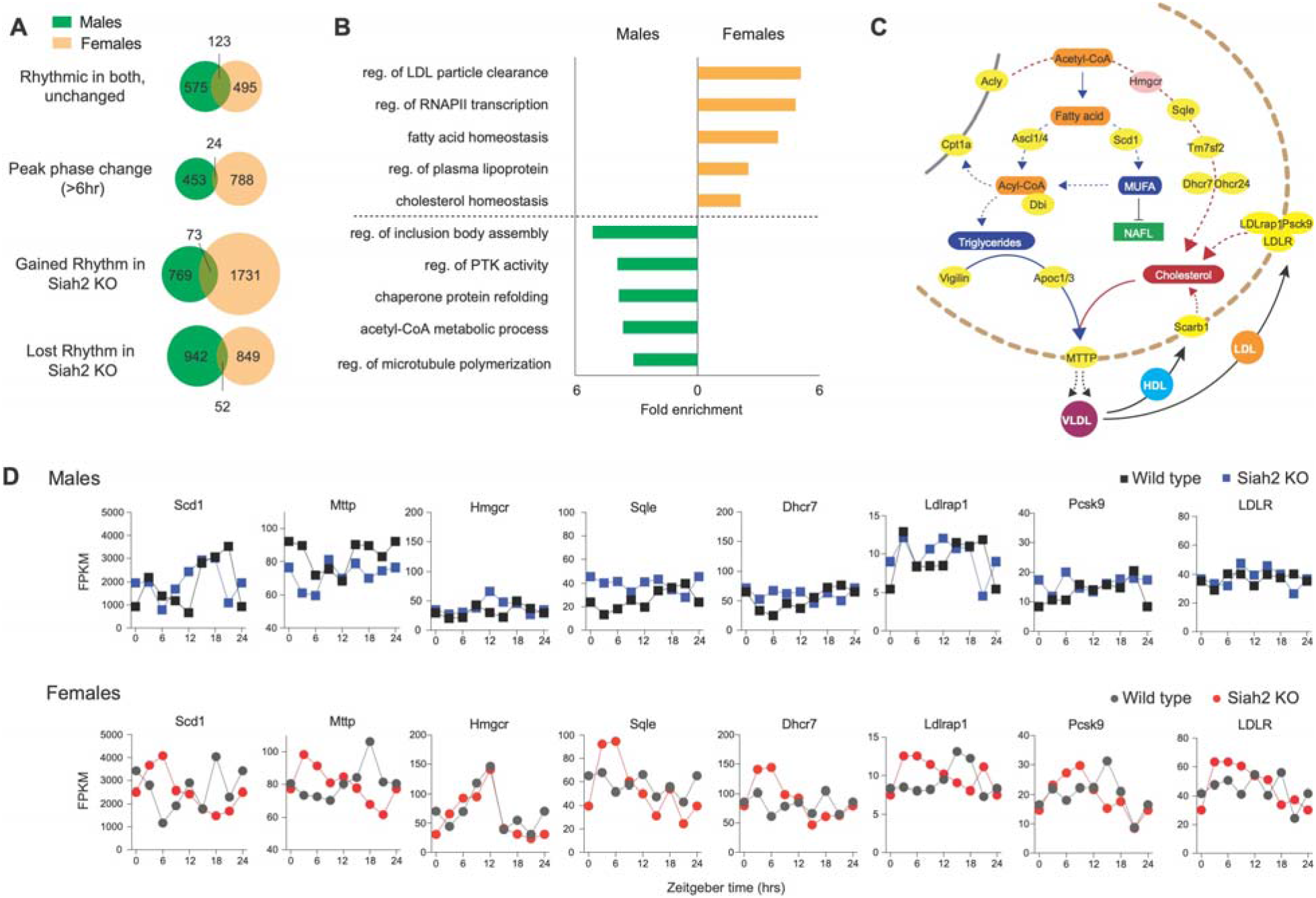
Siah2 regulates rhythmic expression of lipid/lipoprotein metabolism genes in female livers. (**A**) Venn diagrams comparing the effects of Siah2 loss between males and females on rhythmic gene expression. (**B**) Top gene ontogeny terms corresponding to the genes whose expression was affected by Siah2 loss in both sexes. (**C**) Simplified lipid and cholesterol metabolic pathways in the liver, highlighting genes (yellow) uniquely affected by Siah2 loss in females. (**D**) Examples of female-specific effects of Siah2 loss on rhythmic gene expression. See also **Supplemental Datasets 2-3**.

However, before embarking on identifying these mechanisms, we sought to first determine how they may contribute to overall rhythmic physiology. The shift in overall rhythmic gene expression patterns caused by Siah2-deficiency in females creates a prominent misalignment between the timing of gene expression with the normal locomotor and feeding rhythms of *Siah2* KO females (Fig 3F-G). Circadian misalignment between feeding and hepatic gene expression rhythms is known to lead to obesity and metabolic disorders (6, 24–26). Moreover, Siah2 loss in females, but not in males, altered the rhythmic expression of a preponderance of genes involved in regulating gene expression and lipid and lipoprotein metabolism (Fig 4). Thus, the female-specific change in circadian landscape in Siah2-deficient mice led us to predict that these mice would have sex-specific alteration in lipid/lipoprotein regulation and broad deficits in metabolic homeostasis.

We first tested this prediction by asking if the time-dependent changes in the expression of lipid-metabolic genes in *Siah2* KO females (Fig 4) led to alterations in the rhythmic profiles of serum lipoproteins and fatty acids. Siah2 loss in females impacted the rhythmic expression of many key genes invovled in lipid and lipoprotein metabolism (Fig 4C-D), several (i.e. *Mttp, Ldlr, Psck9*) that have direct roles in regulating triglyceride/VLDL and cholesterol levels in the serum, suggesting that any effects of Siah2 loss in females would be restricted to the day. We therefore collected serum at 3-hour intervals across the day from new cohorts of females fed normal chow *ad libitum*. Consistent with the overall change in circadian transcriptional regulation of lipid-regulatory genes, we found that female Siah2 KO mice displayed robust day-time specific increases in serum cholesterol, triglyceride and phospholipid levels (Fig 5A). Moreover, these effects were limited to early daytime and did not appear to disrupt normal nighttime profiles, consistent with the changes in gene expression. In addition, serum free fatty acid (FFA) levels were slightly increased throughout the day and decreased across the night, creating a significant diurnal rhythm in FFA levels in female Siah2 KO mice (Fig 5B). Increases in serum lipids during the day were specific to females (Suppl. Fig 5). Thus, changes to rhythmic gene expression in female *Siah2* KO mice correlate well with the daytime-specific changes in rhythms of lipoprotein/lipid metabolism in the animal, suggesting the two phenotypes are functionally related.

**Fig. 5.**
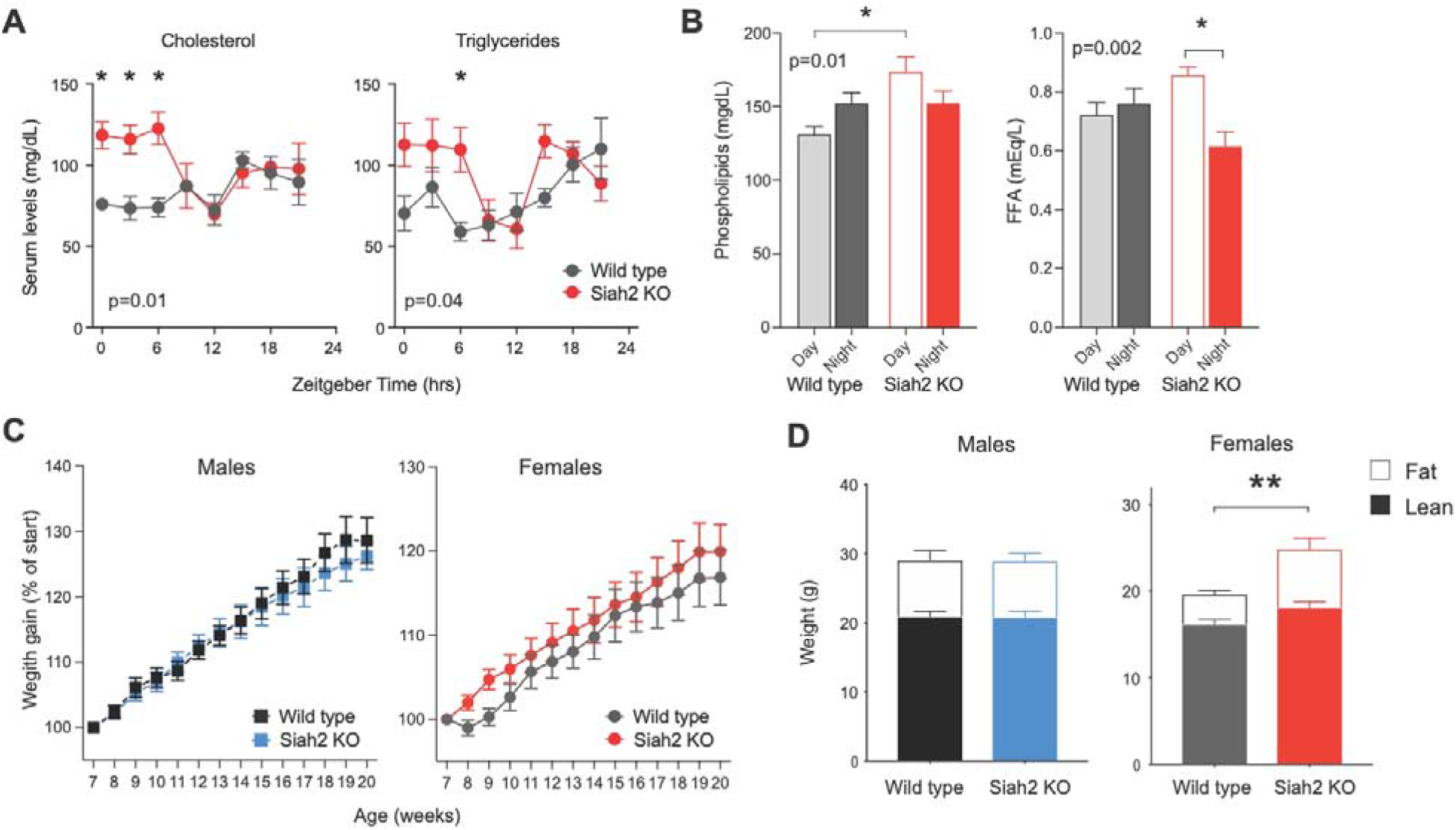
Siah2 loss alters lipoprotein and lipid metabolism in females. (**A-B**) Serum lipid levels across the day in female mice fed normal chow *ad libitum*. In A, each point is the mean +/− SEM (n=3 mice, except ZT 9, n=4). In B, times 0-9 levels were averaged for day (+/− SEM, n = 13 each), times 12-21 averaged for night (+/− SEM, n=12 each). The p-values shown denote significant genotype x time interactions from two-way ANOVA. * = p<0.05 between genotypes (A) or times of day (B) using Sidak’s multiple comparison test. (**C)**. Weight gain in mice fed control diet (CD) Mean +/− SEM, n= 8-10. (**D**) Body composition analysis. Mean +/− sem, n=4-5 mice/sex and genotype. Loss of Siah2 significantly increased adiposity in females on control diet for 13 weeks (** = p=0.0247, two-tailed t-test)

The daytime increase in serum lipids occurs when mice are predominantly sleeping and not eating, functionally similar to eating when the body is sleeping/fasting and can cause obesity (6, 24–26). We tested this as part of the next experiment (see below) by monitoring weight gain and body composition changes through young adulthood. We found that while there was not a detetable difference in overall body weight at 20 weeks of age, Siah2 KO females had a doubling of body fat content compared to wild type females (Fig 5C). Siah2 KO males, in contrast, were not different. Thus, the loss of Siah2 in females, but not males, alters overall fat metabolism, leading increased fat accumulation under normal conditions, presumably as a consequence of the altered rhythmicity of lipid-regulating genes. This also suggests female Siah2 KO mice may be especially sensitive to developing obesity.

To test this possiblity, we metabolically challenged young adult normal and Siah2-deficient mice by switching them to a diet rich in fats. High-fat diets (HFD; 45 kCal from fat) are widely used to exacerbate differences in fat metabolism that underlie development of obesity and other aspects of metabolic syndome, at least in males. It is also well known that, compared to males, females are resistant to the development of HFD-induced obesity and most other metabolic consequences, due at least in part to complex interactions between sex hormones and chromsomes (27–30).

However, we found that female, but not male, *Siah2* KO mice diplayed robust metabolic phenotypes when fed HFD (Fig. 6) compared to wild types. Siah2 KO females displayed a marked increase in diet-induced obesity that was detectable after 7 weeks on HFD (Fig 6A). After 13 weeks, this increased obesity resulted in an overall ~20% weight gain in Siah2 KO mice over wild type females (Fig 6B), despite comparable food consumption (Fig 6C). The weight gain in Siah2-deficient females was predominantly due to an increase in adiposity (Fig 6D), consistent with a deficit in fat metabolism. However, adipocytes of HFD fed Siah2-deficient mice were larger than wild type mice of either sex (Suppl. Fig S6A), suggesting that the sex-specificity in obesity that we observed may not directly involve adipocytes-specific processes (i.e. regulation of PPARgamma; Kilroy et al., 2012). Siah2 also loss altered how HFD impacted the regulation of serum cholesterol and triglyceride levels, selectively in females (Suppl. Fig S6B), and rendered them susceptible to HFD-induced hepatic steatosis (Fig 6E). Again, Siah2 KO males did not show any differences from wild type males in these studies. Unexpectedly, glucose homeostasis remained largely intact in obese Siah2 KO females (Suppl. Fig. S7), suggesting that the mechanistic relationship between diet-induced obesity and diabetes may be sex-specific. Nonetheless, these data suggest that high-fat diet exacerbates an underlying lipid metabolism deficit that is specific to adult Siah2-deficient females. The overall concordance in the sex-specificity of the effects on rhythmic metabolic gene expression and lipid metabolism in mice on normal diets with the female-specificity of consequences of HFD strongly support the notion that the changes in the rhythmic gene expression patterns are likely strong contributors to the metabolic phenotypes in Siah2 deficient female mice.

**Figure 6.**
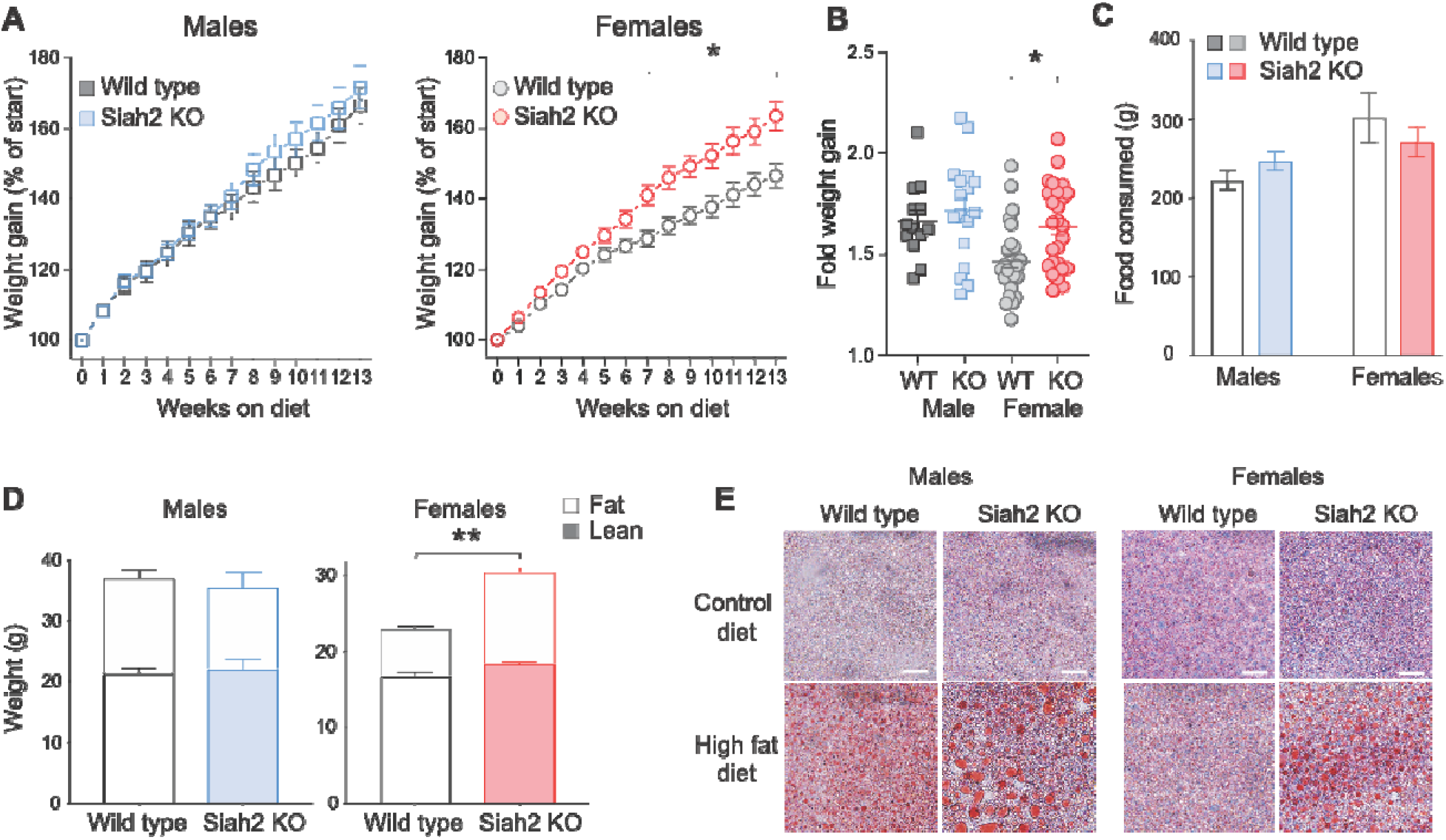
Siah2 protects females from diet-induced metabolic disorders. (**A**) Weight gain in mice fed high fat diet (45% fat). n = 13-17 male mice, combined from two trials, or 26-27 female mice from 4 trials. Two-way ANOVA indicated a significant interaction between genotype x time on diet in females only (p<0.0001, F (13, 663) = 6.484), with significant differences in weight gain by week 7 (* = p <0.01, Sidak’s multiple comparison test). (**B**) Fold weight gained at 13 weeks for the mice in **A**(* p <0.0001, t = 4.809, df = 714). (**C**) Consumption of HFD over the experiment; no statistical differences between any groups. (**D**) Loss of Siah2 significantly increased adiposity in females on HFD for 13 weeks (** p=0.0044, t=4.13, DF=15). (**E**) Oil Red O staining of liver from mice fed HFD for >13 weeks. Red staining = fat. Scale bar = 50 microns.

## Conclusions

Taken together, our data suggest that Siah2 is a key component of an unexpected and unknown female-specific circadian clockwork mechanism that links circadian timekeeping to outputs that regulate rhythms in physiology. Siah2 is a ubiquitin ligase with no known ability to directly regulate gene expression. Thus, Siah2 likely acts on one or more transcription factors that have sexually dimorphic roles in regulating circadian gene expression (Suppl. Fig S4). Identifying these unknown transcription factors is of high interest, as they appear to have strong roles in regulating rhythmic expression of output genes and modulate peak expression of core clock genes in females. It is possible that these transcription factors are female-specific in either expression or function, or how they are intertwined with circadian clock. Indeed, there are >100 transcription factors that appear rhythmically expressed only females (see Fig 3D and Supplemental Dataset 3), but why these are rhythmic only in females and which may be regulated by Siah2 may provide a starting point in identifying these mechansims. At this point, it does not seem likely that this role of SIAH2 is a result of changes in at least global REVERBα and NCOR1 stability (32), and does not appear to involve estrogen or growth hormone signaling. Interestingly, juvenile (4 week old) Siah2 KO mice fed HFD have also been shown to have sex-specific metabolic traits (33) that are somewhat different than those we observed, suggesting possible complex interactions between *Siah2*, diet and puberty. Our data further suggest this is a novel mechanism, as the consequences of Siah2 loss on gene expression selectively target circadian expression patterns and expand, rather than reduce, the overall differences between males and females. In the liver, this mechanism appears to be important for control of lipid metabolism, aligning daily metabolic rhythms in females to their behavior across the day. In this way, this novel Siah2-dependent circadian mechanism may contribute to resilience against diet-induced obesity in females and the overall sexual dimorphism in metabolism. In addition, our findings reveal that circadian clocks can drive gene expression and physiological rhythms using different molecular pathways in males and females. What these mechanisms are, however, still need to be discovered. Nonetheless, the sex-differences in these circadian mechanisms are essential to recognize and decipher as they may contribute why males and females cope differently with circadian clock-related disorders (34–38) and possibly other dimorphisms that exist between males and females.

## Data availability

The full RNAseq data that support the findings of this study are available from the corresponding author upon reasonable request and have been deposited in Gene Expression Omnibus with the accession codes (to be provided prior to publication).

## Acknowledgements

We wish to thank Drs. David Bowtell (Peter MacCallum Cancer Centre) and Andreas Möller (QIMR Berghofer) for providing Siah2 KO breeders. We also want to express strong appreciation to Dr. Zach Hall and members of the DeBruyne, Ehlen and Tosini labs, as well as members of the Neuroscience Institute for valuable feedback and advice for preparing this manuscript. This work was funded by NIH NIGMS grants 1SC1GM109861 and 1R35GM127044 to JPD, as well as in part by NIH NIMHD grants 8G12MD007602, 8U54MD00758, 1G20RR031196, S21MD000101, and C06RR18386 to Morehouse School of Medicine. GT is supported by NIH NEI grant R01EY026291. JCE is also supported by NIH NIGMS grant SC1 1GM127260, JCE, GT and JPD are also supported by NIH NINDS grant U54 NS083932, and JCE and JPD are also supported by IOS NSF grant No. 1832069.

## Supplementary Materials

## Materials and Methods

### Animals and diets

Wild type and *Siah2* KO mice maintained on a C57Bl6 background (1), kindly provided by Dr. Andreas Moller (QIMR Berghofer) and David Bowtell (Peter MacCallum Cancer Institute) were bred and maintained in our animal facility at Morehouse School of Medicine. After an initial cross to C57Bl6 mice, heterozygotes were intercrossed, and homozygous wild type and *Siah2* KO mice were used to establish related lines. All mice had *ad libitum* access to normal chow (PicoLab Laboratory Rodent Diet 5L0D, LabDiet, St. Louis MO, USA) and water and were housed in a 12-hour light:12-hour dark (12:12 LD) cycle unless otherwise stated. Breeding mice were fed breeding chow (PicoLab Mouse Diet 20 5058, LabDiet, St. Louis MO, USA) and all other mice were fed normal rodent chow unless otherwise indicated. For the metabolic studies, mice were individually housed and provided Research Diets D12451 (45% kCal fat) as a high-fat det (HFD) or the sucrose matched control diet (CD) D12450H, starting at 7-8 weeks of age; body weights and food consumption were monitored weekly thereafter. After 13 weeks on these diets, mice were subjected to additional tests (see below). At the indicated times, mice were euthanized using CO_2_ followed by decapitation to collect blood and tissues. All animal studies were approved by the Institutional Animal Care and Use Committee of Morehouse School of Medicine, and in accordance with the United States Public Health Service Policy on Humane Care and Use of Laboratory Animals.

### Body composition and serum analyses

Body composition was performed on mice that were fasted for 6 hours (starting at ZT0, lights off) prior to sacrifice. Body composition was performed post-mortem using MRI at the NIH University of Cincinnati Mouse Metabolic Phenotyping Center (MMPC). Serum was collected following euthanasia with CO_2_ and decapitation and samples were sent for analysis of insulin, total cholesterol, triglyceride, phospholipid and non-esterified fatty acid levels at the same facility. Sex hormones were measured from serum collected single-housed females in proestrus, as determined by cytological examination of vaginal lavages (2) and submitted to Ligand Assay & Analysis Core at the University of Virginia (https://med.virginia.edu/research-in-reproduction/ligand-assay-analysis-core) for measurements of LH/FSH (mouse/rat multiplex), estradiol (mouse/rat) and progesterone (mouse/rat) levels.

### Glucose and insulin tolerance tests

Mice were weighed at ZT0 and then fasted for 4 hours after which the fasting glucose levels were taken. Mice were then injected intraperitoneally with either 1.5mg/gram body weight glucose solution (20% in 0.9% NaCl) or 0.75 IU of insulin per gram body weight (in 0.9% NaCl) and blood glucose levels were measured by drawing blood from the tail 20, 40, and 100 minutes post-injection using a Contour Blood glucose monitoring system (Bayer) as we have done previously(3). Mice were first subjected to glucose tolerance tested, allowed to recover for 1 week before insulin tolerance testing.

### Liver and adipose staining

Oil Red O staining was performed on 16-micron frozen liver sections from wild type or *Siah2* KO, male or female mice fed HFD using the Abcam Oil Red O Kit (ab150678), according to the manufacturer’s instructions. The stained sections were mounted and imaged at a magnification of 40x. Perigonadal white adipose tissue was dissected from the indicated mice, stored in a 15% sucrose at 4°C, and prepared for whole-mount staining(4). Briefly, approximately 4mm x 4mm x 2mm sections of adipose tissue were washed in PBS for 10 minutes then stained sequentially with DAPI (Thermofisher; D1306) and Cell Mask Orange (ThermoFisher; C10045) for 1 hour each, with PBS washes between. Images and measurements were obtained using a spinning disc confocal microscopy system and Slidebook 6 (Intelligent Imaging Innovation, Denver CO, USA).

### RNA isolation and quantitative real-time PCR

Liver samples from *Siah2* KO and wild type mice were collected every 3 hours over 24 hours (n=3 per genotype and time point). Total RNA was extracted from livers using Trizol reagent (Invitrogen) per manufacturer’s instructions. Approximately one microgram aliquots of total RNA were reverse transcribed and subjected to quantitative PCR using Applied Biosystems cDNA kit (Thermo Fisher) and SsoAdvanced SYBR GREEN qPCR mix (Bio-Rad) respectively. Samples were run using the CFX96 Touch Real-Time PCR Detection System (running version 3.1 of the CFX Manager Software; Bio-Rad) and data all normalized to Gapdh and plotted relative to the time-independent average of the wild type samples using the 2^−ΔΔCt^ method. The sequences of the primers used were as follows: m*GAPDH*: forward: AGACAGCCGCATCTTCTTGT, reverse: CTTGCCGTGGGTAGAGTCAT; mNr1d1: forward: CCCTGGACTCCAATAACAACACA, reverse: GCCATTGGAGCTGTCACTGTAG; *mArntl*: forward: AACCTTCCCGCAGCTAACAG, reverse: AGTCCTCTTTGGGCCACCTT; *mPer2:* forward: GAAAGCTGTCACCACCATAGA, reverse: AACTCGCACTTCCTTTTCAGG; *mDbp:* forward: GGAACTGAAGCCTCAACCAAT, reverse: CTCCGGCTCCAGTACTTCTCA; *mCry1:* forward: TGAGGCAAGCAGACTGAATATTG, reverse: CCTCTGTACCGGGAAAGCTG.

### Transcript profiling

We performed two different transcript profiling experiments using slightly different approaches. For the data shown in Fig 1, livers were collected at ZT10 from mice of both sexes and genotypes at ~20 weeks of age, and total RNA was extracted using Trizol, followed by clean up using RNeasy kits (Qiagen). Transcripts were profiled using the Lexogen QuantSeq 3’ mRNA kit to ‘count’ transcript abundance. Library prep and sequencing was performed by Omega Bioservices (Norcross, GA, USA). Data were trimmed, mapped and analyzed using Illumina’s BaseSpace platform. Differential expression was determined using RNAexpress and the DESeq2 method. Full count data and DESeq2 results are provided in Supplemental Dataset 1. Enrichment of rhythmic genes in these data was determined using the “Mouse 1.0ST Liver” and “Mouse Liver 48 hour Hughes 2009” datasets (http://circadb.hogeneschlab.org/mouse), with a probabilty cut-off value at q<0.05 using the gene symbols for all 513 differentially expressed genes as search terms.

For the diurnal transcriptomics, livers were isolated from *Siah2* KO and wild type mice maintained on normal chow at 3-hour intervals across a 12:12 LD cycle. Total RNA was isolated and cleaned-up as described above, and equal amounts were pooled from livers from 3 mice/time/genotype/sex for RNAseq. RNAs were converted into sequencing libraries by using Illumina TruSeq stranded mRNA Library Prep kits and sequenced by Omega BioServices (Norcross, GA) using the Illumina HiSeqX10 platform. Samples were sequenced to a depth > 37 million 150bp X 150bp paired end reads. The reads were mapped to the mouse MGSCv39-mm9 genome using Tophat 2.1.0. Expression levels were assessed using Cufflinks 2.2.1, which calculates raw counts and the number of fragments per kilobase per million (FPKM). Low expression genes were filtered out if the sum of their raw count was less than 100 across the time points in all 4 groups (male wild type, male *Siah2* KO, female wild type and female *Siah2* KO). Count data was used for DeSeq2 (https://yanli.shinyapps.io/DEApp/) (Fig. S2), and FPKM was used for all other analyses.

### Time Series Analysis for Circadian Cycling

MetaCycle::meta2d (ver.1.2.0; https://CRAN.R-project.org/package=MetaCycle) was used to detect circadian transcripts with the default settings and period length set to 20 for ‘minper’ and 28 for ‘maxper’(5). Data from one cycle was concatenated to create a 48 hour time series for analyses using JTK cycle(6) to identify cycling transcripts and their peak expression phases and DODR ver.0.99.2(7) (https://CRAN.R-project.org/package=DODR) to directly compare gene expression profiles between sexes/genotypes. We acknowledge that concatenating the data like this likely increases the false-positive rates in both algorithms. However it also greatly reduces the false-negative rate that especially JTK can have when analysing a single cycle ; JTK was developed to operate more effectively with two cycles of data. False negatives for rhythmicity are much more problematic for our analyses as these would likely falsely amplify the differences between groups – something we sought to minimize as much as possible. To help limit the overall effect of false-positives, we combined results from 2-3 measures (i.e. rhythmicity, phase, meta.p) from both analyses to define differences (described below). It should be noted that performing these analyses on non-concatenated data produces the same overall proportionality in the results – *Siah2* KO females still have ~60% more rhythmically expressed genes than wild type at all cut-off levels (Fig S1A), with similar changes in phase distribution (Fig S1B-C), but with fewer numbers of genes in each category. The peak times (i.e. phase) were derived using JTK cycle and imported into Oriana (version 3, Kovach Computing, Anglesey, Wales UK) to produce Raleigh plots and GraphPad Prism (v7 or later; GraphPad Software, SanDiego CA, USA) for further analyses. Heat maps were generated using an R code (https://github.com/gangwug/SRBR_SMTSAworkshop/blob/master/R/fig.R).

The goal of JTK-cycle is to identify rhythmic transcripts within a dataset but does not perform direct comparisons between datasets. DODR identifies overall differences between time-series datasets, including phase, amplitude and overall abundance, but does not assess circadian-like rhythmicity per se, thus can identify changes even in transcripts that are not rhythmic in either dataset being compared. Therefore, we used JTK cycle parameters to define rhythmicity and peak phase, and DODR to substantiate differences/lack of differences between groups, as depicted in the table below (bold font indicates key difference).

|  | Group 1 JTK<br>adjP value | Group 2 JTK<br>adjP value | DODR meta.p | Phase<br>difference |
| --- | --- | --- | --- | --- |
| Rhythmic in both,<br>unchanged | < 0.05 | < 0.1 | < 0.01 | **< 6 hours** |
| Rhythmic in both,<br>phase change | < 0.05 | < 0.1 | Not used | **> 6 hours** |
| Rhythmic in both,<br>other difference | < 0.05 | < 0.1 | **> 0.01** | < 6 hours |
| Gained Rhythm/<br>Females only | < 0.05 | **> 0.1** | **> 0.01** | < 6 hours |
| Lost Rhythm/<br>Males only | **> 0.1** | < 0.05 | **> 0.01** | < 6 hours |

We chose these JTK adjP value cutoffs in attempt prevent overestimating differences between groups, while still including genes with less robust rhythms. Using the same cutoffs for both groups increases the number of different genes between groups, but reducing the cutoffs proportionally reduces the numbers of genes in each category, but the differences between groups remain proportional (i.e. there are still more differences in females than males). DODR was not used to support phase changes, but meta.p > 0.01 for 85% of genes with Siah2-loss induced phases differences in females (Fig 4a), 75% of those changes in males (Fig 4a) and 82% of sexually dimorphic phase changes (Suppl. Fig S3F) genes (**see Supplemental datasets 2-4**). In addition, we found small subsets of genes we classified as rhythmic in across both groups, but DODR identified a difference other than a 6-hour phase change were not closely examined but are denoted in **Supplemental datasets 2-4**. Of note, the differences in gene expression observed via qRT-PCR (Fig 1) were not readily detectable in the RNAseq data due to sample pooling and the differences in analysis methodologies. All of the differentially expressed genes were combined and subjected to Gene Ontology analyses (http://geneontology.org). Top relevant child terms for biological processes were selected after sorting by decreasing order of fold enrichment and increasing order of p-values.

### Locomotor and feeding behavior

Wheel running locomotor activity was recorded and analyzed as described previously (8). For feeding behavior, mice were fed ad-libitum with normal chow and eating behavior was recorded for 8 days using infrared video cameras (ZOSI 720p CVI TVI, ZosiTech, Zhuhai city, China) connected to a TigerSecu 8 Channel DVR Security Video Recording System. Videos were analyzed using Noldus EthoVision XT (v14, Noldus, Leesburg VA, USA), and feeding behavior was coded in 1 min bins using the following criteria: 1) the mouse took food from the feeder with its mouth or, 2) the mouse moved food in the feeder with its mouth; either behavior having persisted for 3 seconds or more. When eating behavior occurred, that 1 min bin was coded as “1”. One-minute bins without any feeding behavior were coded as “0”. One-minute bins were summed for each hour and averaged across the 8-day recording according to ZT hour in 1-hour intervals for each mouse, and then normalized to the within animal mean of its total daily feeding activity.

### Statistical Analyses

Except where noted above, all graphs and statistical analyses were generated using Graphpad Prism (v7 or later; GraphPad Software, SanDiego CA, USA). Statistical analyses performed were typically two-way ANOVAs, examining *sex* x *genotype* or *genotype* x *diet* interactions, followed by Sidak’s multiple comparison test, unless otherwise indicated. Relevant F values and degrees of freedom are also reported as ‘F (DFn, DFd) = [value]’ or t=[value], df=[value]. Differences were considered significant if p<0.05, unless otherwise indicated.

## Datasets S1-S5. (separate files)

**Supplemental dataset 1** lists the QuantSeq results for all transcripts detected. Official gene symbol, counts for each individual, means for each genotype and the results of Deseq2 analysis. Data are sorted by *padj* value. The differential genes in ‘Circadian Rhythm’ DAVID term are highlighted in red font. **Supplemental datasets 2-4** list the genes/NM identifiers and FPKM data, and the JTK-cycle and DODR results from comparing wildtype and *Siah2* KO females (**Dataset 2**), wild type and *Siah2* KO males (**Dataset 3**) or wild type males and wild type females (**Dataset 4**); a ‘key’ for the column labels is provided in each. These data are parsed according to difference within the comparison (described below). **Dataset 5** lists the genes with time-independent changes in gene expression, due to either Siah2 loss (in males and females) or sex. DeSeq2 results are included. The full RNAseq datasets have been deposited in NCBI GEO, under the identifiers (provided prior to publication).

**Figure S1:**
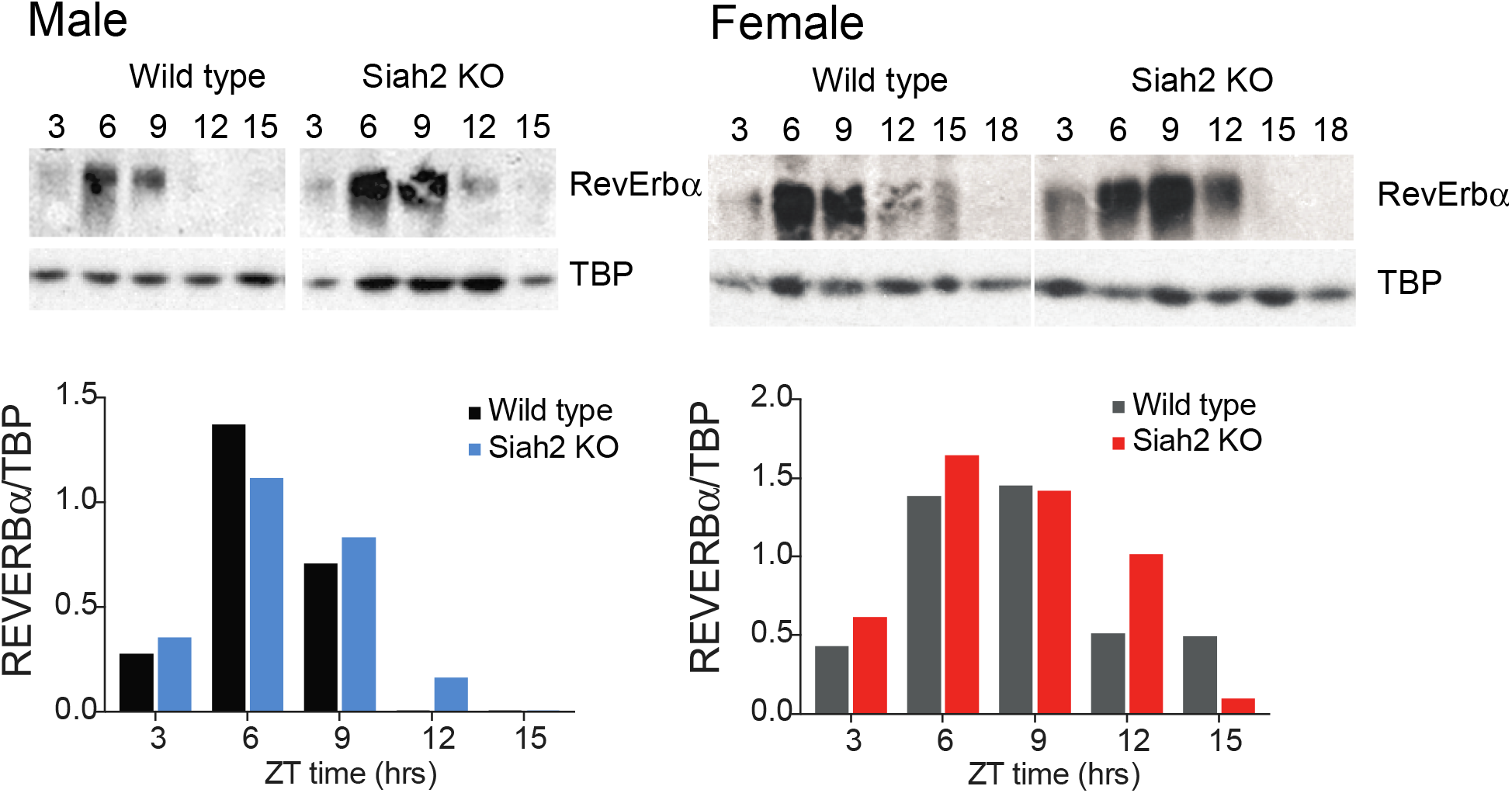
Siah2 loss does not drastically alters REV-ERB α stability in livers. Representative western blots of pooled liver samples (n=3 mice/pool) collected around the clock (time 12 = lights out). Blots are shown above their quantitation using ImageJ.

**Fig. S2.**
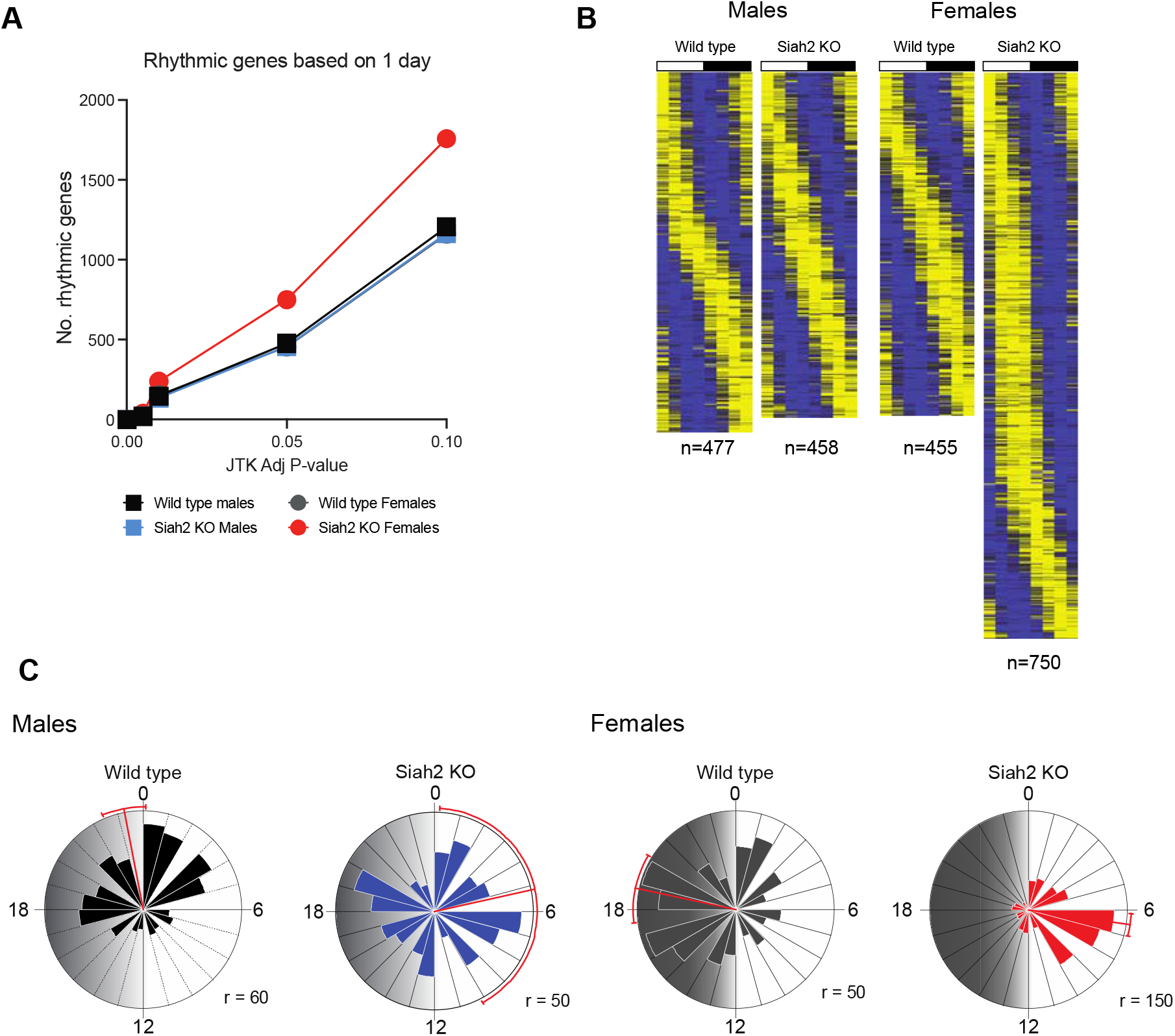
Analysis of single-day data profiles. **A**. Plot of the number of genes are rhythmic based on different JTK cutoffs in each group. Wild type males and females, as well as Siah2 KO males all have similar numbers of rhythmic genes across cutoffs; Siah2 KO females have ~50-60% more rhythmically expressed genes at any cutoff. **B**. Heatmaps of the genes rhythmic at AdjP <0.05 for each group, plotted independently. **C**. Frequency distribution of peak expression timing for the genes plotted in B. Numbers around the clock face are hours, ZT time (light and dark are indicated by shading). Red lines and error bars are the mean peak timing +/− 95% CI for the entire population, r = radii of the circular plots in no. of genes.

**Fig. S3.**
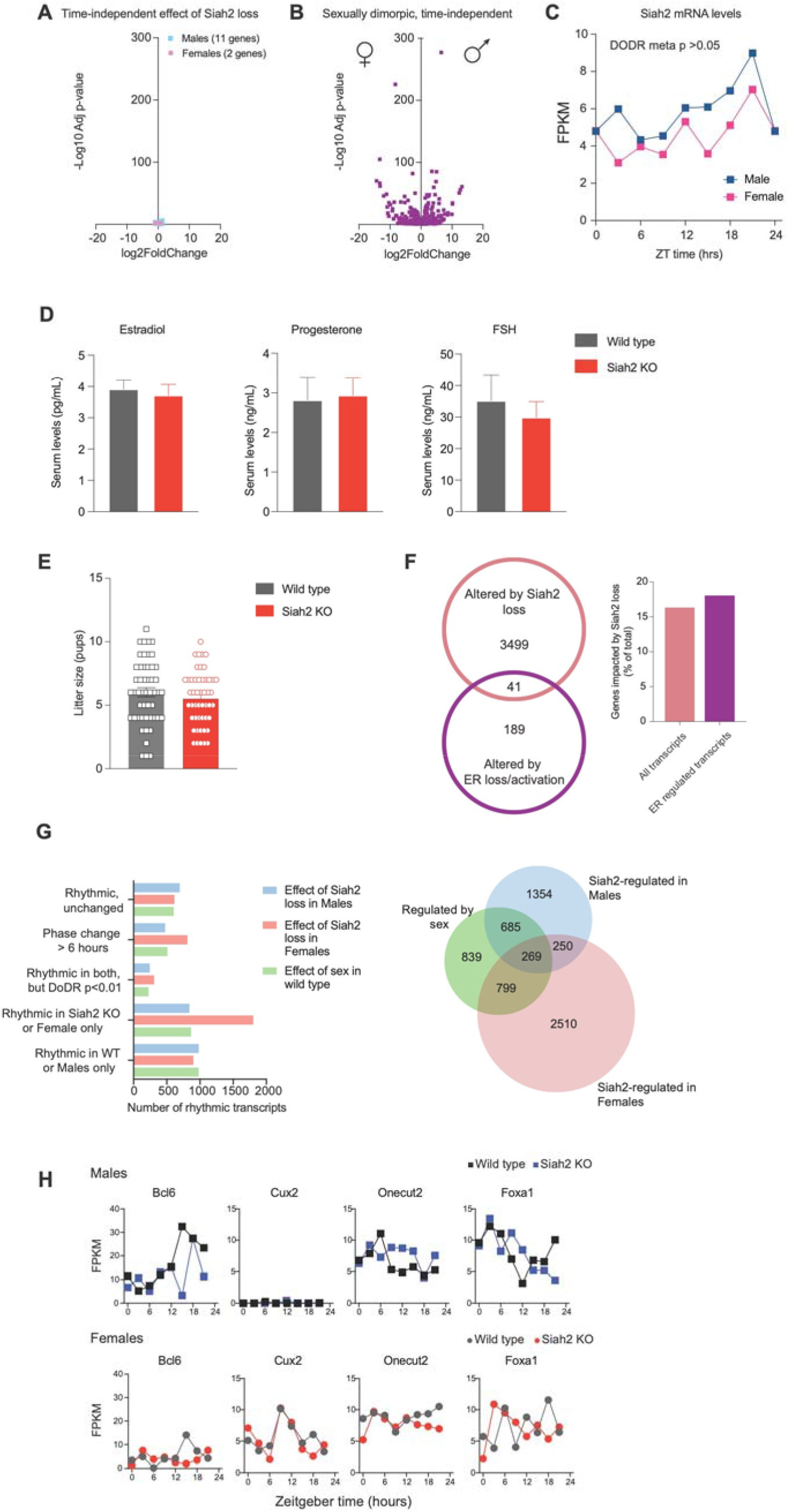
Siah2 loss does not drastically alter time-independent gene expression or preferentially target common sexually dimorphic pathways. **(A)** RNAseq data were combined across timepoints according to sex/genotype, treating timepoints as biological replicates (n=8 per sex/genotype) and the effects on Siah2 loss on gene expression in each sex were compared using DeSeq2. This volcano plot shows the few genes that were significantly different (pAdj <0.05), which are plotted scale identical to those shown in **B** for comparison. **(B)** Time-independent gene expression comparison between wild type males and females, as described for A. Only the significantly different (pAdj <0.05) transcripts are shown for clarity. The sex symbols indicate higher expression in female or male livers, respectively. Genes for both A and B are listed in **Supplemental Dataset 5. (C)** Siah2 expression itself is not sexually dimorphic. RNAseq FPKM of Siah2 in wild type male and female livers. Overall there was no detectable difference in expression (DoDR meta p >0.05, DESeq2 padj>0.05). **(D)** We collected serum from females in proestrus sacrificed between ZT 7-14 for measurement of the indicated hormones (LH levels were below level of detection). Mean +/− SEM (n=7 each, except n= 5 for wild type progesterone) hormone levels are shown. p > 0.5 for each, unpaired two-tailed t-tests. (**E)** Litter sizes for last 50/49 litters obtained from 18/19 homozygous wildtype/homozygous *Siah2* KO females (respectively), bred with males of like genotypes. Individual data points and mean +/− SEM are shown (p = 0.3, t=1.038, df=97, two-tailed t-test). **(F).** *Left-* Venn diagram depicting the overlap between genes regulated by Siah2 loss and estrogen signaling in female livers (9). *Right* - Proportions of transcripts whose expression was altered by Siah2 loss in female livers. Siah2 loss altered the expression of 3,540/22,002 total genes, or ca. 16% of the total genes examined. Siah2 loss altered a similar proportion (ca. 18%) of estrogen responsive genes suggesting that Siah2 loss is not selectively impacting estrogen-signaling. Thus, the effects of Siah2 loss on rhythms is unlikely to occur via mis-regulation of estrogen signaling. **(G)** Comparison of the effects on diurnal gene expression of Siah2 loss in males and females and the effect of sex in wild type mice, categorized as described in the **Supplemental Materials and Methods**. Some of these data are replotted from **Figure 2** for comparison. See also **Supplemental Datasets 2-4**. Venn diagrams depicting the overlap in genes with dimorphic rhythmic expression with those differentially expressed by Siah2 loss in male or female livers, across the categories in **C**. See also **Supplemental Datasets 2-4. (H)** RNAseq profiles for representative genes encoding factors involved in mediating sexual-dimorphic gene expression in the liver.

**Fig. S4.**
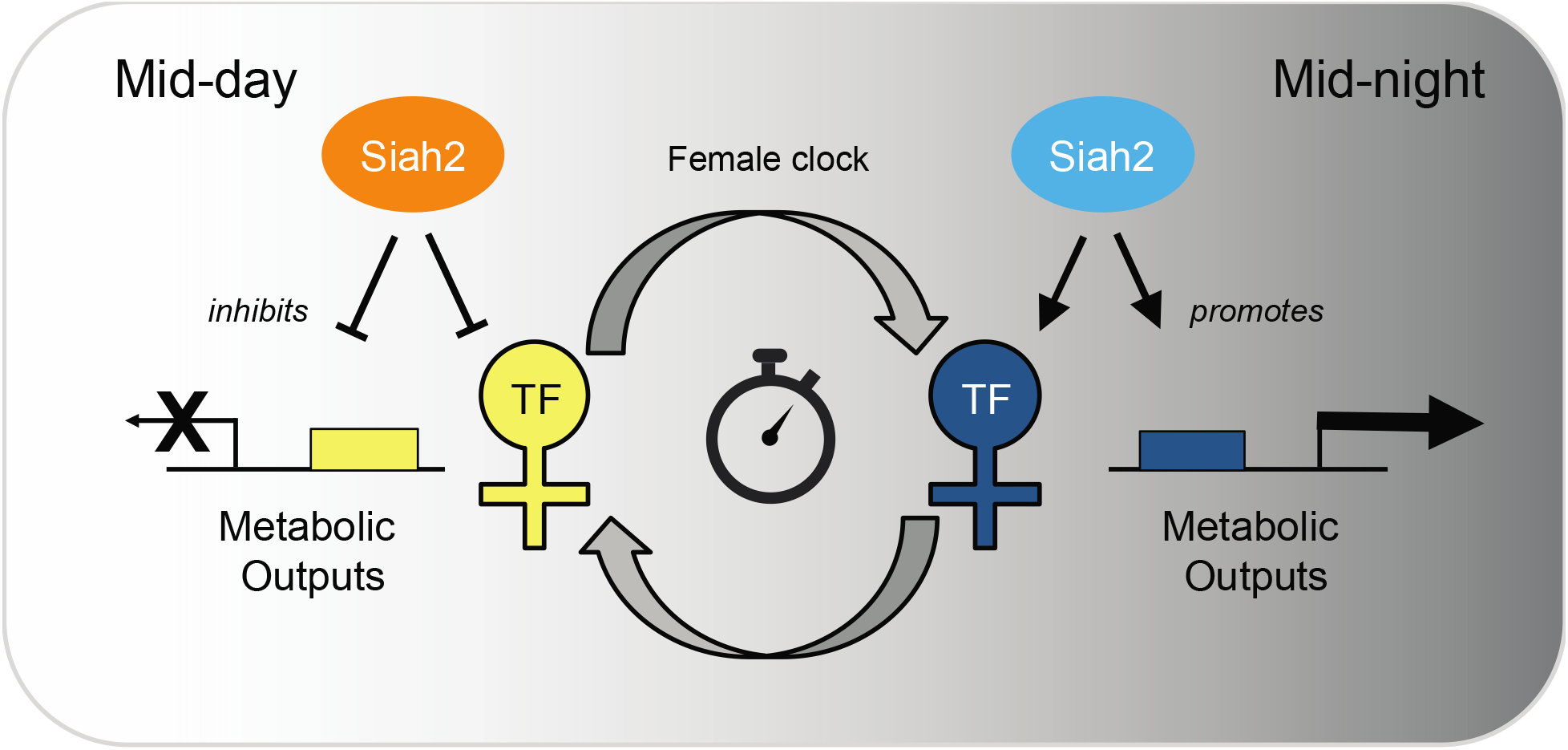
Model depicting possible intersecting points of Siah2 with the female circadian clockwork. Siah2 appears to have opposing roles on regulating rhythmic gene expression in the female liver. Our data suggest Siah2 plays a strong role in repressing gene expression at mid-day, while also promoting expression of many genes during the night. In addition, for some targets, Siah2 appears to advance/delay normal nocturnal expression to the daytime. Since Siah2 is not known to directly bind DNA, we propose that Siah2 is mediating these effects via regulating unknown transcription factors that play a female-specific role in overall circadian regulation.

**Fig. S5.**
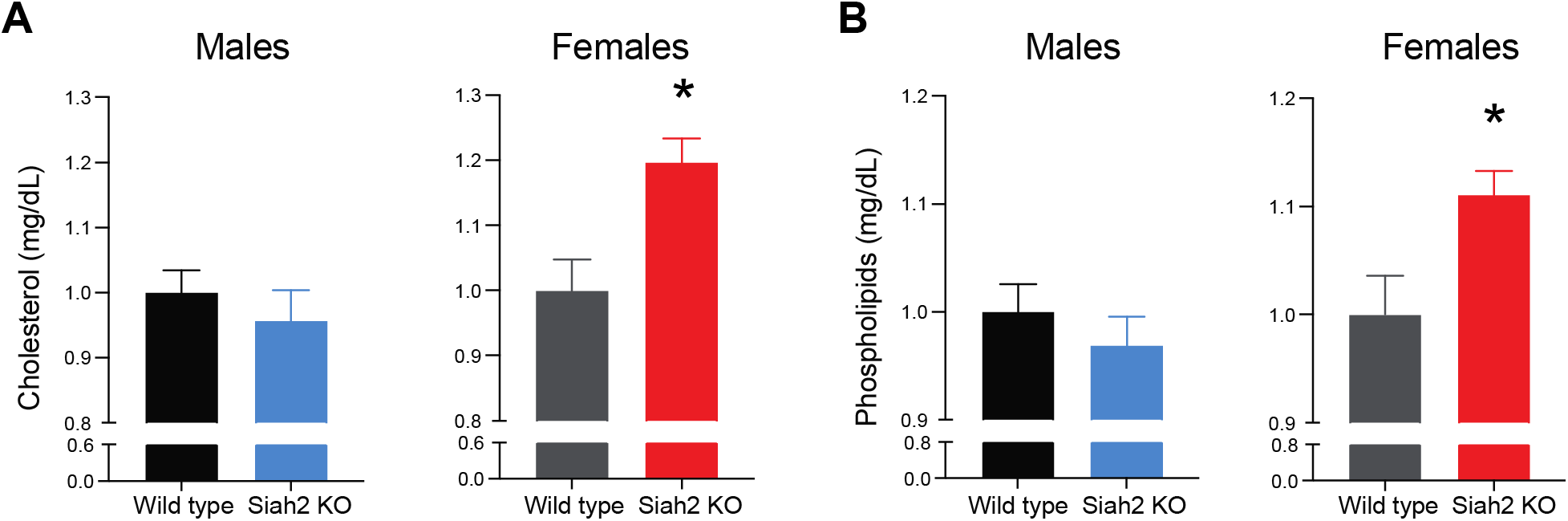
Effects of Siah2 loss on serum lipoproteins are female specific. Serum was harvested from mice sacrificed between ZT3-6 and assayed for total cholesterol (A) and phospholipids (B). Data are the mean +/− SEM, n=15 combined from three independent trials. Data are normalized to wild type to eliminate trial-to-trial differences overall levels. * = p <0.05, two way t-test from wild type.

**Fig. S6.**
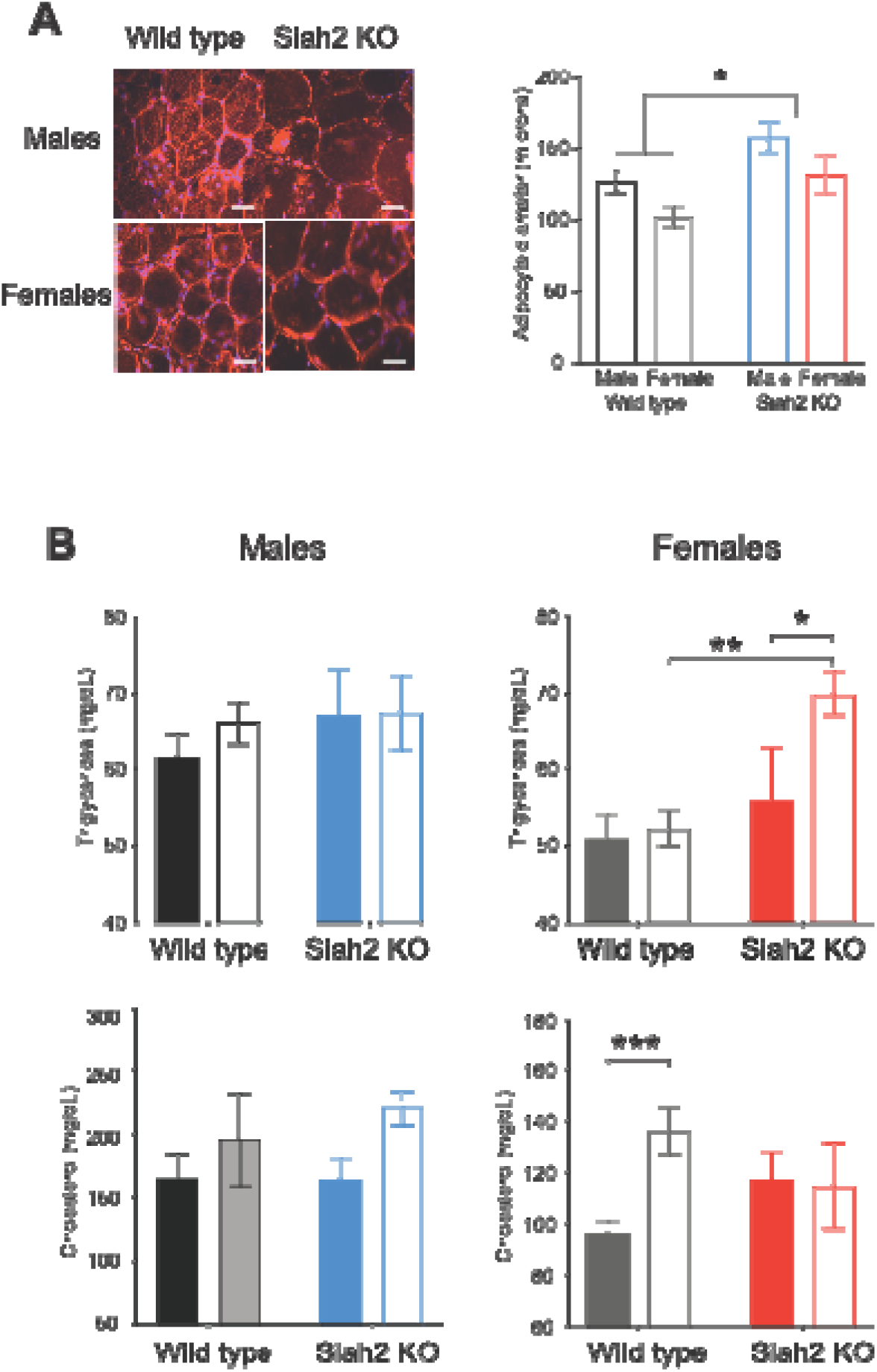
Effects of HFD on adipocyte size and serum lipid regulation in Siah2 KO mice. **(A)** Left – Representative images of adipose samples obtained from mice fed HFD for >13 weeks and stained with Cell Mask Orange (red) and DAPI (blue) to visualize cell membranes and nuclei, respectively. Scale bar = 50 microns. Right – quantitation of 100 cell diameters measured per mouse (mean +/− sem, n=3 mice/group). * p=0.018 (F (1, 8) = 8.836), main effect of genotype, two-way ANOVA. **(B)** Serum triglycerides (upper) or total cholesterol (lower) in response to 13 weeks of feeding CD (dark bars) or HFD (light bars). Sample were collected at ZT10 after a 6 hour fast. Mean +/− SEM are shown (n=5 mice sampled per group, except n= 4 for Siah2 KO males or females fed CD). * p = 0.048 (t=2.05, df = 15), ** p = 0.009 (t=30342, df = 15), *** p = 0.043 (t=2.563, df=15), Sidak’s multiple comparisons test.

**Fig. S7.**
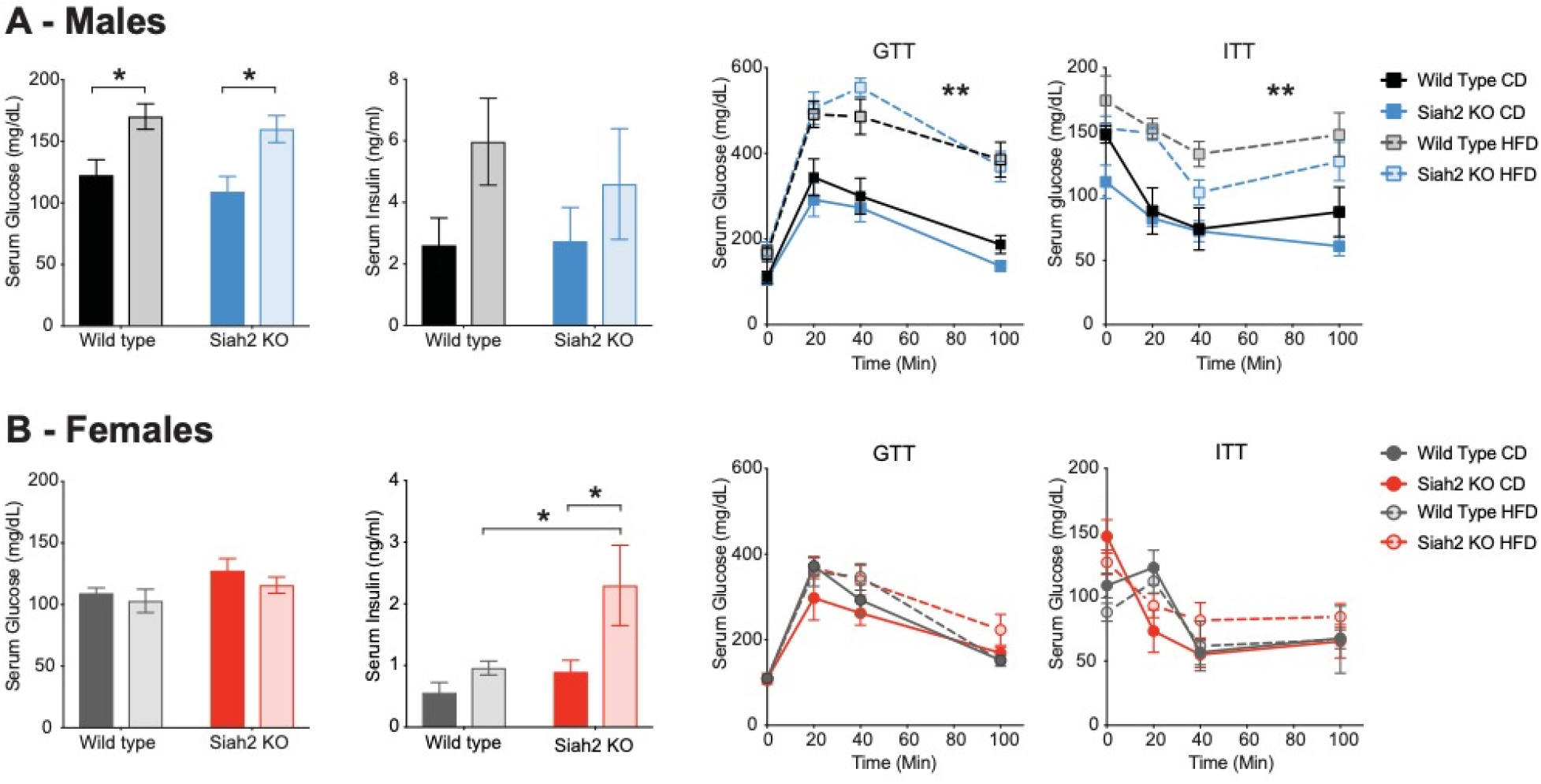
Obese *Siah2* KO females do not show strong diabetic-like phenotypes. Fasting serum glucose and insulin levels, as well as glucose and insulin tolerance (GTT and ITT, respectively) in males **(A)**and females **(B)**. All animals were fasted for 4 hours (starting at ZT0) before testing. Data are all means +/− sem, n=5-7 mice per group, except n = 4 for wild type CD males and wild type HFD females. * = p<0.03, ** p=0.0009 for time x diet interaction (F (3, 72) = 6.140), three-way ANOVA), and main effect of diet (p<0.0001, F (1, 72) = 121.0) but no three-way interaction (p=0.79, F (3, 72) = 0.3480), *** main effects of diet (p<0.0001, F (1, 72) = 68.48”) and genotype (p=0.0043, F (1, 72) = 8.688) only, no three-way interaction (p=0.6376, F (3, 72) = 0.5684). ^#^ p=0.039, t=2.605, df=15 and ^##^ p=0.037, t=2.64, df=15 using Sidak’s multiple comparison test.

